# Forward reactivation in human hippocampal CA1 supports repetition learning and is preserved in the aging brain

**DOI:** 10.64898/2026.10.20.760541

**Authors:** Zhiyao Gao, Li Zheng, Vinod Menon, Weidong Cai

## Abstract

Repetition is a fundamental mechanism for strengthening episodic memory, yet the neural computations engaged during re-exposure remain disputed. Competing accounts attribute repetition-related gains either to retrospective reinstatement of prior events or to the prospective use of learned structure to anticipate what comes next. Using high-resolution functional MRI and representational similarity analysis in a lifespan cohort (N = 418, ages 36–100), we tracked hippocampal representations as participants twice encoded two fixed sequences of face–name pairs, resolving reactivation by direction, distance and sequence. During repeated encoding, CA1 and CA2/3/DG representations spontaneously anticipated immediate next states, unprompted by explicit task demands. Forward, but not backward, reactivation predicted repetition-related gains, and showed no detectable age dependence. Only CA1 forward reactivation predicted memory gains after accounting for reactivation in other medial temporal lobe regions. Repetition learning is thus an active, temporally local prospective computation with a distinctive CA1 contribution that appears preserved well into old age.

## Introduction

Repetition is among the oldest and most reliable means of strengthening memory ^1^. From the oral transmission of knowledge in preliterate societies to modern educational curricula and clinical rehabilitation, repeated exposure remains the most widely used method by which new information is converted into durable knowledge. Controlled studies have established across materials, timescales, and populations that repeated encounters improve retention ^2–4^. What repetition does, computationally, remains far less clear. A second encounter with information is not a duplicate of the first: the learner arrives carrying a trace of the original episode, so the brain state that reencodes the stimulus has itself been changed by prior experience. Characterizing what the brain does with that prior trace is central to understanding why repetition works and why repetition may benefit older adults even as other forms of learning decline.

Any account of repetition learning is fundamentally a claim about memory traces, about what happens to the representation of an event when the event recurs. Early neuroimaging studies of repetition learning focused on regional activation, establishing that hippocampal and neocortical responses are reliably modulated by repeated presentation ^5–8^. Such measures demonstrate that repetition changes processing, but they cannot indicate which representation is active at a given moment, and so cannot distinguish a strengthened copy of the original trace from one that has been reorganized or has become predictive. Multivariate approaches, which read out distributed patterns of activity rather than their amplitude, make that distinction tractable and are critical for probing the mechanisms of repetition-based learning ^9,10^.

Contemporary theories propose that the benefits of repetition arise through study-phase retrieval: on re-encountering an item, the brain reactivates the representation formed during the first encounter, and the mnemonic benefit follows from that reactivation ^11–13^. Whereas classical work examined reactivation during offline states, such as sleep or rest ^14–20^, emerging evidence suggests that reinstatement can also occur dynamically during active encoding ^11,21^.

A central unresolved question concerns the temporal direction of neural reactivation, with two competing accounts ^22^. The backward reactivation account holds that repeated exposure reinstates the prior encounter in order to stabilize and strengthen the original memory trace, functioning as a retrospective reinforcement mechanism ^22^. Predictive-processing theories instead hold that the brain is fundamentally anticipatory, and that repetition allows prior experience to be used to forecast upcoming stimuli and optimize learning ^23–25^. Evidence for distinct temporal functions comes largely from rodent work showing that forward replay is linked to consolidation and planning whereas reverse replay is more closely associated with reward learning ^26–28^. Consistent with a prospective bias in humans, behavioral studies report that forward associations may form stronger memory bonds than backward associations in temporal sequences ^29–31^, and forward reinstatement has been observed in the human hippocampus during naturalistic movie viewing ^32^ and statistical learning ^33^. It therefore remains unclear which direction predominates when repetition confers a measurable memory benefit. It is equally unknown how these processes relate to the strength of the initial trace: strong initial encoding might provide the substrate for better subsequent prediction, or anticipatory processing might instead be recruited preferentially for traces that were initially weak.

The medial temporal lobe (MTL), and the hippocampus in particular, is central to repetition learning ^34,35^. Beyond indexing past experience, the hippocampus is increasingly understood as a predictive map encoding expected future states ^25,36–38^. This framework makes predictive reactivation computationally advantageous for repetition learning: repeated experience may allow hippocampal networks to forecast upcoming stimuli rather than merely reinforce prior traces ^35^. The hippocampus is not itself a unitary structure but is comprised of specialized subfields with different memory representations ^39–42^, and temporally precise prediction should depend on coordinated computation across them. Classical rodent replay studies implicate CA1 and CA3 place cells ^22^; whether analogous subfield-specific mechanisms support episodic repetition learning in humans is an open question. The temporal scope of any such predictions is also undefined: human hippocampal forward reactivation might span entire sequences or operate locally, forecasting only the immediate next state.

Episodic memory declines across mid-to-late adulthood (Grady & Craik, 2000)^43,44^, accompanied by age-related changes in MTL structure and function and reduced fidelity of hippocampal representations ^42,45–48^. Nevertheless, older adults continue to benefit from repeated encoding ^49–51^, and recent evidence indicates comparable repetition-learning rates in younger and older adults for sequentially presented memory lists but worse repetition learning for pairwise associations ^52^. This relative preservation raises the possibility that reactivation-based computations supporting sequential repetition remain available across age despite MTL vulnerability. Whether forward reactivation provides an age-resilient mechanism through which repeated experience continues to strengthen memory across adulthood remains unknown.

To address these questions, we leveraged high-resolution functional MRI and trial-level representational similarity analysis (RSA) ^53^ to a large lifespan cohort from the Human Connectome Project in Aging (HCP-A; N = 418, ages 36–100) ^54^. Participants performed a face-name sequence memory task featuring repeated learning and recall (**Fig. 1a**). Crucially, because the task used a fixed stimulus order that was identical across repetitions, the serial position of each item provides a common index across exposures, allowing the direction and distance of representational change between the first and second encounter to be quantified directly (**Fig. 1b–e**).

**Fig. 1.**
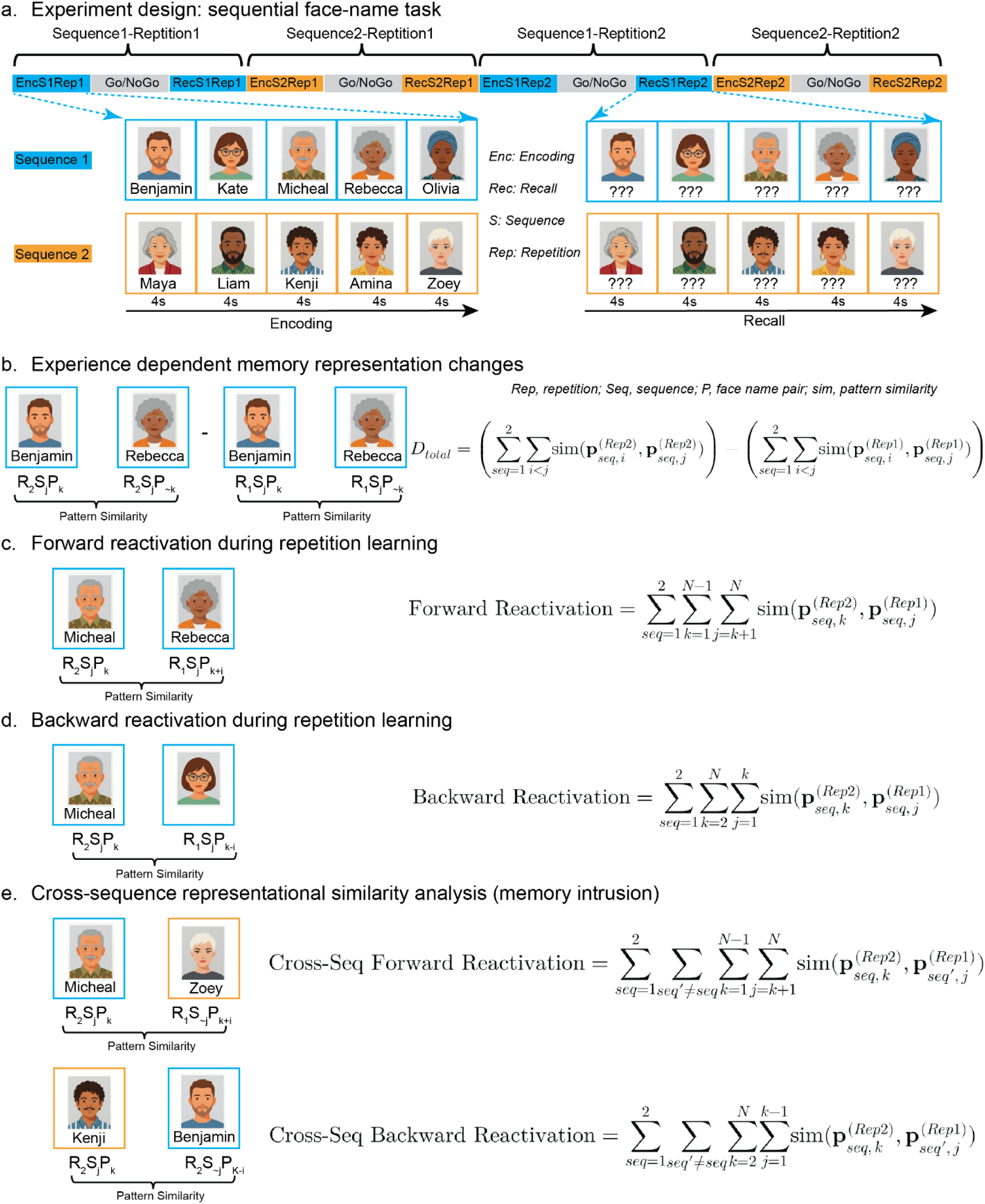
a. Experimental paradigm and analytical framework for sequence memory reactivation. (a) Face-name sequence learning paradigm. Participants completed a fixed-sequence associative memory task featuring two learning repetitions (Rep1 and Rep2). Each repetition comprised an encoding block (memorizing face-name pairs), an active distractor block (a Go/No-Go task to prevent continuous maintenance or rehearsal), and a cued recall block. b. Experience-dependent representational changes. Neural pattern similarity between all stimulus pairs within a sequence was computed for both Rep1 and Rep2. c. Directional memory reactivation analysis, forward reactivation during repetition learning. To disentangle the temporal mechanisms driving memory improvement, experience-dependent representational changes were parsed directionally. Forward prediction was quantified as the neural similarity between a given stimulus during Rep2 and subsequent upcoming sequence items from Rep1. d. Backward reactivation was quantified as the similarity between the Rep2 stimulus and preceding sequence items from Rep1. e. Cross-sequence representational similarity analysis (memory intrusion). Cross-sequence forward reactivation was quantified as the similarity between Rep2 stimulus (k) and stimulus after the order k from the other sequence Rep1. Cross-sequence backward reactivation was quantified as the similarity between Rep2 stimulus (k) and stimulus preceding the order k from the other sequence Rep1. These neural measures were subsequently correlated with behavioral performance to explore whether the benefits of repeated sequence learning, short- and long-term memory are preferentially driven by anticipatory forecasting or retrospective rehearsal.

We first tested whether experience-dependent hippocampal transformations track behavioral memory gains, and parsed their temporal direction to arbitrate between anticipatory forward reactivation and retrospective backward reactivation. We then examined how this relationship varies with initial memory strength and with sequential distance, and whether it is separable from general working memory. Finally, we examined whether the neural correlates of repetition-related memory gains identified in these analyses varied with age.

Our findings provide novel evidence that repetition learning is accompanied by anticipatory hippocampal coding. During re-exposure, representations in CA1 and CA2/3/DG came to resemble the item that was about to appear, and the degree of that resemblance tracked how much memory improved. Our results recast repetition not as passive re-exposure but as an active process in which the hippocampus uses prior experience to represent the immediately upcoming item. This anticipatory signal showed no detectable age dependence, suggesting preserved precision of local prediction may contribute to aging resilience.

## Results

### Demographic and behavioral results

The HCP-A included 680 participants with neuroimaging and behavioral data of the face–name task. After screening (see details in Methods), the final sample comprised 418 participants (56.13±13.47, 236 female, **Fig. 2a**).

**Fig. 2.**
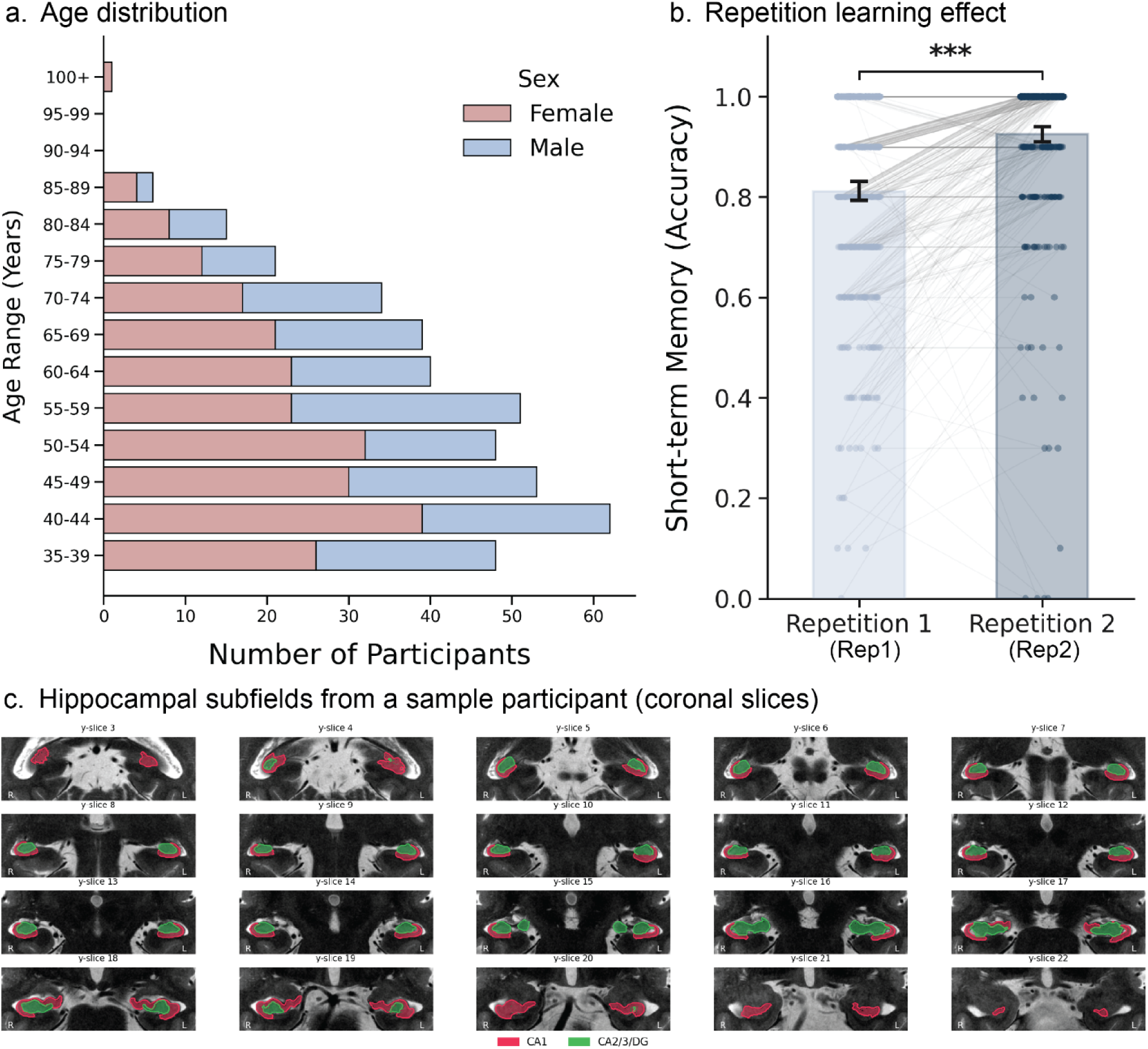
Sample demographics, learning effects, and hippocampus segmentation. (a) Age distribution of the full sample. (b) Repetition learning effect. Participants demonstrated significant increases in short-term memory accuracy over repetition learning. (c) Example segmentation of CA1 and CA2/3/DG in one participant.

Participants showed significant improvements in memory performance with repeated learning. Accuracy increased and reaction time (RT) decreased from the first repetition (Rep1) to the second repetition (Rep2) (*ps* < 0.001, **Fig. 2b**). For subsequent analyses, we defined the repetition learning effect as the within-participant increase in accuracy from Rep1 to Rep2 (Rep2 − Rep1), such that larger values indicated greater repetition-related improvement.

Age was negatively associated with short-term memory performance at both Rep1 (r = −0.139, p = 0.005) and Rep2 (r = −0.218, p < 0.001), long-term memory performance measured outside the scanner (r = −0.356, p < 0.001), as well as MoCA scores (r = −0.163, p < 0.001). In contrast, age was not associated with the repetition learning effect (p>0.2).

Together, these findings suggest that the capacity to strengthen memory through repetition is relatively resilient to age, even as overall memory and global cognition decline.

### MTL subfields volume in relation to age, cognition and memory

Because our primary analyses quantified multivoxel pattern similarity within individually segmented MTL regions, age-related differences in ROI size could affect the number and reliability of voxels contributing to similarity estimates and thereby confound across-participant brain–behavior associations. Accordingly, before testing our main hypotheses, we first characterized structural variation of MTL subfields in gray matter volume and their relationship to cognition and memory. All regional volumes were normalized by intracranial volume to control global brain size.

Normalized volume showed trend-level associations with repetition learning effect in CA1, CA2/3/DG, and the whole HIPP (r = 0.109∼0.123, ps = 0.063, FDR-corrected) after controlling for age, sex, and education. No significant association between MTL volumes and memory performance was found after adjusting for the same covariates. See details in **Supplementary Results**.

MTL subfields exhibited robust age-related volume reductions that remained significant after controlling for sex and education (r = -0.197 ∼ -0.480, all ps < 0.001, FDR-corrected), but not associated with by MoCA (all ps > 0.8).

Together, these results demonstrate a robust age-related structural decline within MTL and weak associations between MTL volume and memory measures. Consequently, normalized volume was included as an additional covariate in all subsequent analyses to account for potential structural confounding.

### Hippocampal representation changes during encoding predict learning effects

We first examined whether the HIPP and its subfields, CA1 and CA2/3/DG, exhibited neural reactivation during Rep2 encoding and whether this reactivation related to memory improvements. These regions were selected a priori because of their established roles in memory reactivation, integration, pattern separation and completion, and the mitigation of memory interference ^35,39,42,55^. Unlike the initial Rep1 encoding, where participants had no knowledge of upcoming stimuli, the repeated Rep2 encoding allowed prediction of subsequent items in the fixed face–name sequences. We hypothesized that within-sequence neural pattern similarity would be greater during Rep2 than Rep1, reflecting predictive memory reactivation, and that these changes would correlate with repetition-related memory gains.

To test this, we performed a region-of-interest (ROI)-based representational similarity analysis (RSA) to quantify neural pattern similarity among all within-sequence stimulus pairs during Rep1 and Rep2 encoding phases. The difference in mean pattern similarity between Rep2 and Rep1 served as an index of memory prediction (**Fig. 1b**). Because Rep1 and Rep2 followed identical trial orders and structures, these effects are unlikely to reflect temporal autocorrelation or perceptual differences. We observed significant positive correlations between repetition-related changes in pattern similarity and repetition learning effect in CA1 (r = 0.164, p = 0.001), CA2/3/DG (r = 0.113, p = 0.023), and the whole HIPP (r = 0.169, p = 0.001), all p values FDR-corrected. These findings remained significant after adjusting for age, sex, years of education, regional volume, head motion, and scanner sites (CA1: r = 0.162, p = 0.004; CA2/3/DG: r = 0.103, p = 0.0498; HIPP: r = 0.159, p = 0.004; all p values FDR-corrected. **Fig. 3a**), indicating that experience-dependent changes in neural representations robustly predict memory improvement. Similar reactivation patterns were found in other task phases (see **Supplementary Results**).

**Fig. 3.**
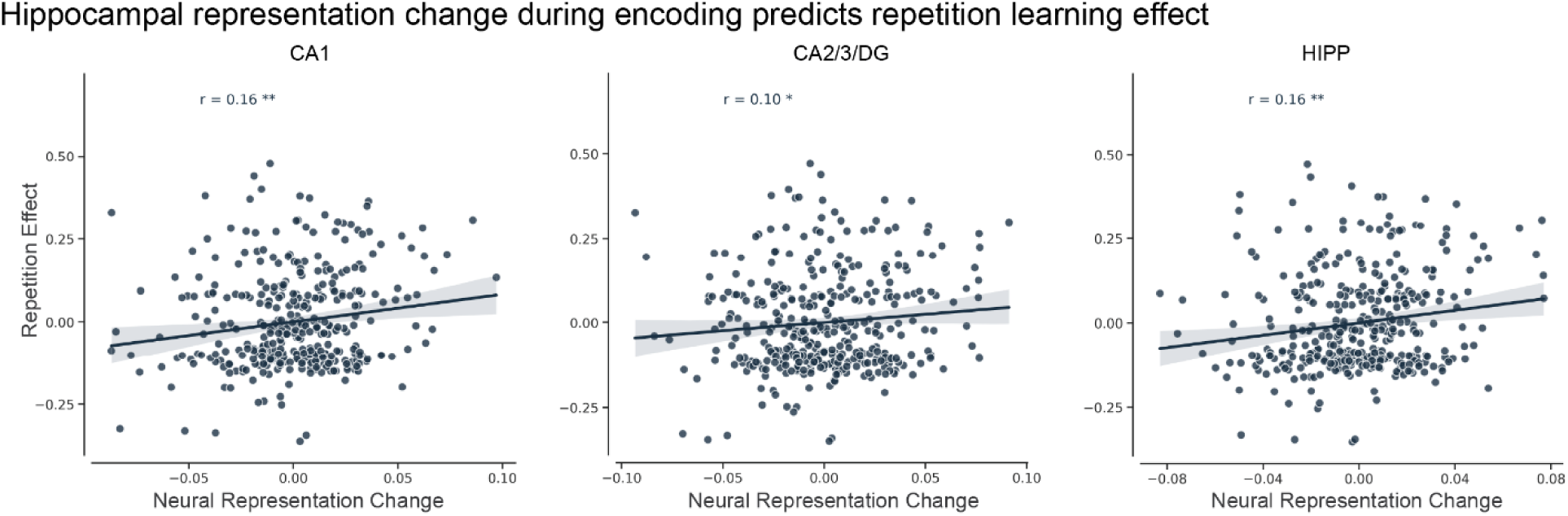
Hippocampal representation change during encoding predicts repetition learning effects (CA1: r=0.16, p = 0.004; CA2/3/DG: r = 0.10, p = 0.049; HIPP: r = 0.16, p = 0.004). For visualization, representation change and repetition learning were each regressed separately on age, sex, education history, head motion, scanner site, and normalized regional volume. The resulting residuals are plotted on the x- and y-axes, respectively. ** p < 0.01; * p<0.05. All p values were FDR corrected.

Additional control analyses examining cross-sequence representation changes in relation to repetition learning effects yielded no significant results across ROIs (ps = 0.732), with all p values FDR-corrected (see **Supplementary Method/Results**), suggesting repetition learning–dependent representation changes being specific to the current repeated sequence.

We next tested whether repetition-related representation change varied with chronological age after adjusting for sex, years of education, normalized regional volume, head motion, and scanner site. No significant age association was observed in any ROI (ps > 0.354, FDR-corrected), indicating no detectable age dependence of the representation change.

### Forward reactivation in the hippocampus predicts repeated learning effect

As the changes in neural representation during Rep2 encoding might reflect rehearsal of previously presented stimuli or prediction of upcoming stimuli, we examined whether brain–behavior relationships were better explained by neural measures indexing forward or backward reactivation. The orders of Face–Name Pairs were identical in Rep1 and Rep2, allowing us to examine backward and forward reactivation by comparing neural representations of items across repetitions according to their relative sequence positions. For example, during Rep2 encoding, stronger forward reactivation would be reflected by greater similarity between Face–name Pair 3 (Rep2P3) and subsequent Rep1 items (Rep1P4 or Rep1P5) (**Fig. 1c**), whereas stronger rehearsal would be reflected by greater similarity with preceding Rep1 items (Rep1P1 or Rep1P2) (**Fig. 1d**).

We found a significant positive relationship between forward reactivation during Rep2 encoding (relative to Rep1) and repetition effect in the CA1, CA2/3/DG, and the whole HIPP (CA1: r = 0.119, p = 0.017; CA2/3/DG: r = 0.132, p = 0.013; HIPP, r = 0.130, p = 0.013; FDR-corrected). The effects remained significant after adjusting for confounding factors such as age, sex, education, regional volume, head motion, and scanner sites (CA1: r = 0.126, p = 0.022; CA2/3/DG: r = 0.121, p = 0.022; HIPP, r = 0.128, p = 0.022; FDR-corrected. **Fig. 4**). In contrast, no significant relationships were observed for backward reactivation/rehearsal before or after adjustment for covariates (ps > 0.7). Similar patterns were observed during other task phases (see **Supplementary Results**).

**Fig. 4.**
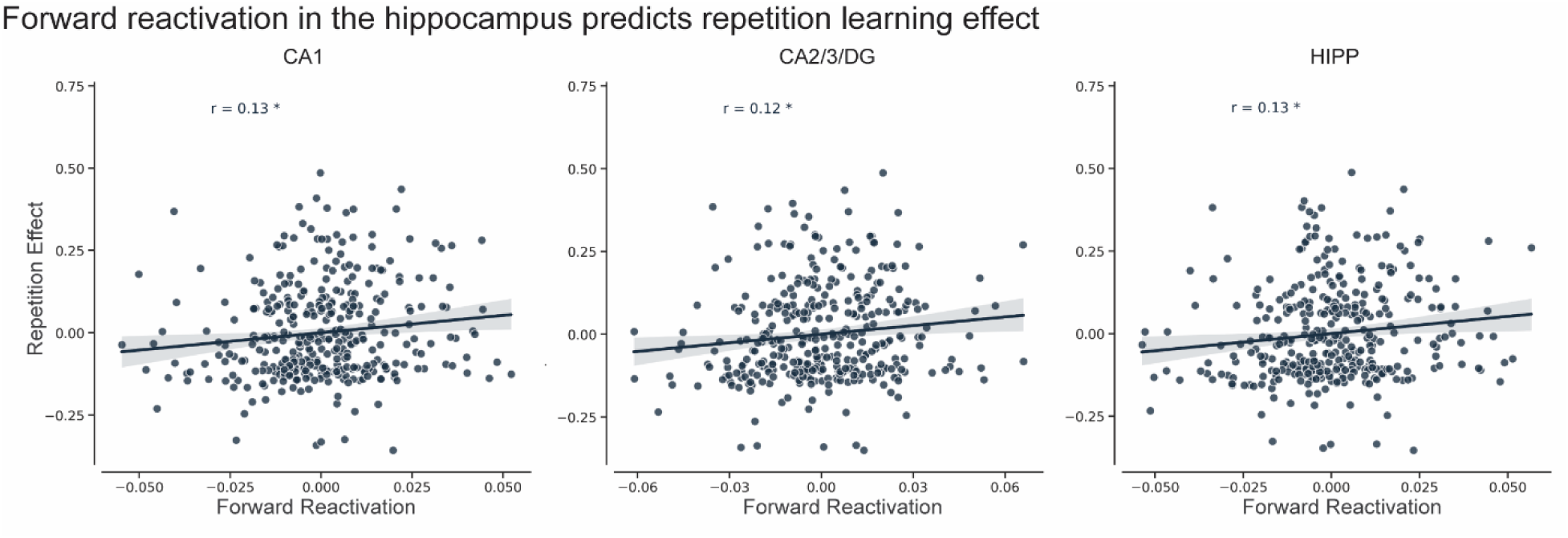
Forward reactivation predicts repetition learning effects. Forward reactivation: CA1: r = 0.13, p = 0.026; CA2/3/DG: r = 0.12, p = 0.022; HIPP: r = 0.13, p = 0.022. Forward reactivation and corresponding behavior were each regressed separately on age, sex, education history, head motion, scanner site, and normalized regional volume. The resulting residuals are plotted on the x- and y-axes, respectively. All p values were FDR-corrected. * p < 0.05.

To test whether these effects reflected memorizing the location rather than specific content, we conducted a control analysis using cross-sequence comparisons (e.g., comparing Rep2P3 from sequence 1 with P4/P5 or P1/P2 from sequence 2, see **Fig. 1e**). No significant brain–behavior relationships were observed in either direction before or after adjustment (all ps>0.3, see details in **Supplementary Results**), supporting the content specificity of the effect.

An alternative explanation is that the forward-similarity measure reflects nonspecific temporal carryover within a repetition rather than prediction. If activity elicited by Rep1 item k persists into Rep1 item k+1, residual item-k information could inflate the similarity between Rep1 item k+1 and Rep2 item k. Critically, the same process should also inflate backward similarity: persistence of R2 item k−1 into Rep2 item k would increase the similarity between Rep2 item k and Rep1 item k−1. A nonspecific carryover account should therefore produce comparable forward and backward brain–behavior associations. Moreover, because this account depends partly on the cross-repetition reinstatement of the same-item representation, we examined whether cross-repetition same-item similarity predicted memory improvement. No such association was observed in any ROI after covariate adjustment (ps = 0.985, FDR-corrected). Together, these findings argue against nonspecific temporal carryover as the primary explanation for the forward association.

To establish the directional specificity of this relationship, we also examined whether forward and backward reactivation explained unique variance in repetition-related memory improvement. In analyses controlling for the standard covariates and the complementary directional measure, forward reactivation remained positively associated with memory improvement in CA1, CA2/3/DG, and the whole HIPP (r = 0.122/0.109/0.132; FDR-corrected p = 0.031/0.040/0.031, respectively). Conversely, backward reactivation was not associated with memory improvement after accounting for forward reactivation (all FDR-corrected ps > 0.727). As a complementary analysis, we computed a lag-controlled directionality index by subtracting backward reactivation from forward prediction. This analysis yielded a convergent positive association for forward prediction in CA1 (see **Supplementary Results**).

Lastly, we also tested whether repetition-related forward reactivation varied with chronological age after adjusting for sex, years of education, normalized regional volume, head motion, and scanner site. No significant age association was observed in any ROI (all FDR-corrected ps > 0.294), indicating no detectable age dependence of the forward-reactivation signal.

Together, these findings provide consistent evidence that forward, but not backward, hippocampal reactivation predicts memory improvement during repeated learning, even when such prediction is entirely incidental to task demands.

### Hippocampal forward reactivation in relation to first-exposure memory

Our previous analyses demonstrated that experience-dependent changes in hippocampal representations during repeated learning were primarily driven by forward prediction during Rep2 encoding. However, it remains unclear whether these predictive reactivation processes depend on the strength of the initial memory trace formed during first exposure (Rep1).

Short-term memory performance during Rep1 was negatively associated with forward reactivation during Rep2 encoding defined by the similarity to neural activation patterns during Rep1 encoding in CA1, CA2/3/DG, and the whole HIPP (r = −0.149/−0.142/−0.136, ps = 0.01; FDR-corrected), after controlling for standard covariates including age, sex, education, normalized volume, head motion, and scanner site (**Fig. 5a**). Crucially, this effect remained significant after accounting for working memory as an additional covariate (ps = 0.032, FDR-corrected), demonstrating its independence from working memory capacity.

**Fig. 5.**
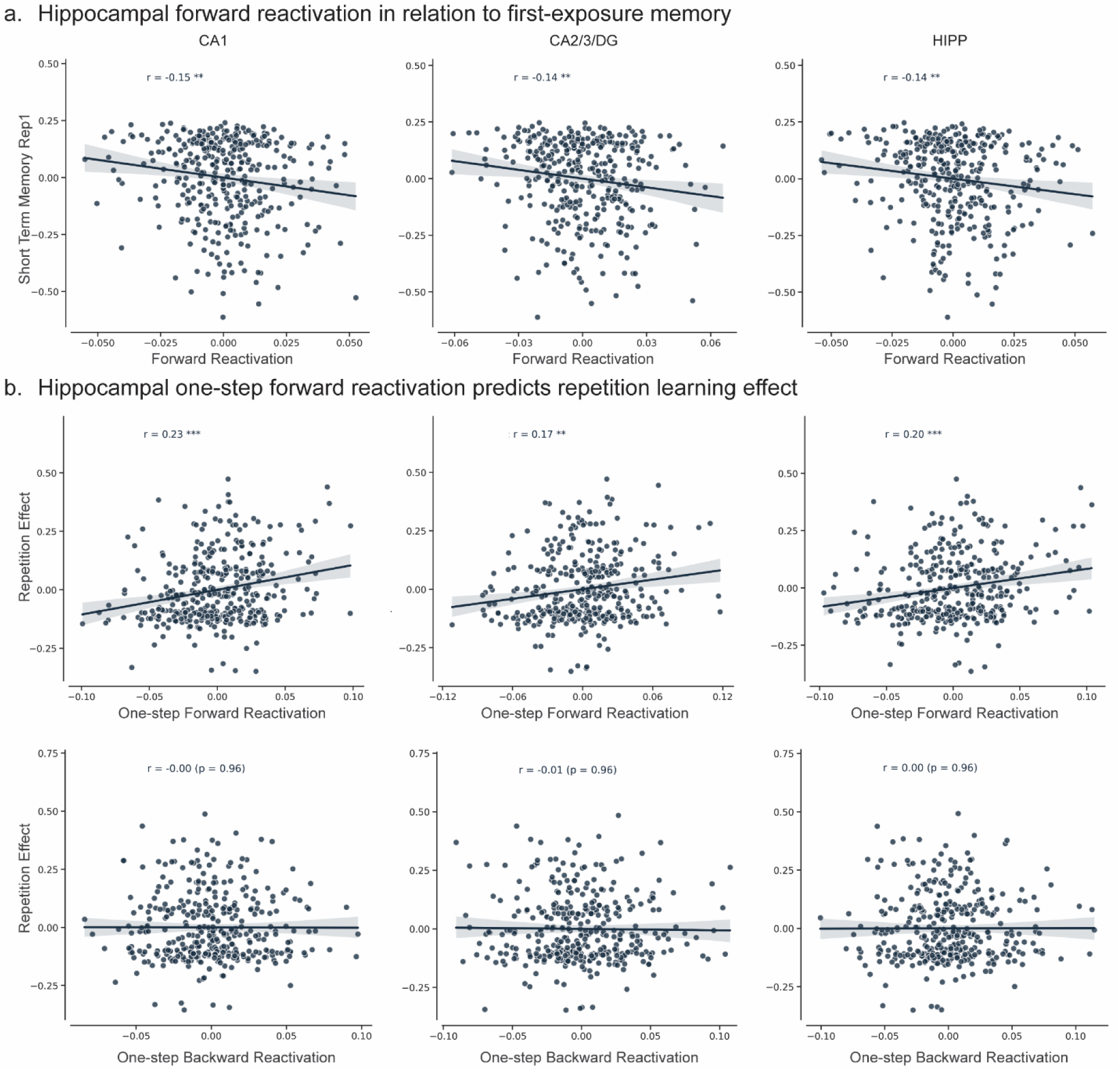
a. Initial weaker short-term memory was associated with stronger forward reactivation during repeated encoding (CA1: r = -0.15; CA2/3/DG: r = -0.14; HIPP: r = - 0.14; ps = 0.01). b. One-step forward reactivation predicts repetition learning effects (CA1: r = 0.23, p < 0.001; CA2/3/DG: r = 0.17, p = 0.001; HIPP: r = 0.20, p < 0.001). Forward/backward reactivation and corresponding behavior were each regressed separately on age, sex, education history, head motion, scanner site, and normalized regional volume. The resulting residuals are plotted on the x- and y-axes, respectively. A 10,000-resample bootstrap analysis showed that one-step forward reactivation was more strongly associated with repetition learning than one-step backward reactivation in CA1, CA2/3/DG, and HIPP (p = 0.012/0.024/0.012). All p values were FDR-corrected. *** p < 0.001; ** p < 0.01; * p < 0.05. All p values were FDR-corrected.

Collectively, these findings indicate that forward reactivation during the second exposure of information is inversely related to short-term memory of the first exposure. It is not merely a byproduct of generalized cognitive limitations—such as poor working memory impairing initial Rep1 encoding or restricting the resources available to predict future representations. Rather, this suggests that predictive hippocampal reactivation may serve as a compensatory function during repeated relearning by strengthening initially weaker memories.

### Hippocampal forward reactivation in short- and long-distance forecasts

Next, we examined how far ahead memory representations anticipate upcoming information, and which predictive distances best support repetition learning. Because aggregate measures may obscure temporally specific effects, we separately analyzed discrete predictive distances (e.g., one-step vs. two-step forward prediction). This approach also allowed us to test whether the absence of backward reactivation effects in prior analyses masked localized backward signals.

We observed significant positive associations between one-step forward reactivation during Rep2 encoding when similarity was compared to Rep1 encoding and repetition-related learning effects in CA1, CA2/3/DG, and the whole HIPP (r = 0.228/0.167/0.201; p = 0.0000/0.0014/0.0002; FDR-corrected), after controlling for standard covariates (**Fig. 5b**). In contrast, no significant relationships were found for any other forward or backward reactivation distances (ps > 0.125, FDR corrected). A 10,000-resample bootstrap analysis showed that one-step forward reactivation was more strongly associated with repetition learning than one-step backward reactivation in CA1, CA2/3/DG, and HIPP (p = 0.012/0.024/0.012, FDR-corrected).

Moreover, after adding one-step backward reactivation to the standard covariates, one-step forward reactivation remained positively associated with repetition learning in CA1, CA2/3/DG, and HIPP (r = 0.223/0.175/0.203, respectively; all FDR-corrected p < 0.001). In the reverse analysis, one-step backward reactivation was not significantly associated with repetition learning in any ROI after adjustment for one-step forward reactivation (FDR-corrected ps = 0.959). We also calculated a directionality measure by subtracting one-step backward from one-step forward reactivation. After adjustment for the standard covariates, this measure was still positively associated with repetition learning in CA1, CA2/3/DG, and HIPP (r = 0.159, 0.118, and 0.152; FDR-corrected p = 0.006, 0.025, and 0.006, respectively). Finally, to assess potential carryover effects, we added cross-repetition within-item pattern similarity to the standard covariates. One-step forward reactivation remained positively associated with repetition learning in CA1, CA2/3/DG, and HIPP (r = 0.230, 0.171, and 0.211, respectively; all FDR-corrected p ≤ 0.001).

Finally, we examined if this one-step forward reactivation was associated with chronological age after adjusting for sex, years of education, normalized regional volume, head motion, and scanner site. No significant age association was observed in any ROI (all FDR-corrected ps > 0.762). Thus, the precise one-step-ahead representational signal linked to repetition-related memory strengthening showed no detectable age dependence across the sampled age range.

### Unique contribution of CA1 forward reactivation to repetition learning

To determine whether the association between hippocampal reactivation and repetition learning reflects a distributed hippocampal mechanism or is driven by specific subregions, we tested whether whole-hippocampus one-step forward reactivation still predicted repetition learning after adding reactivation in each subregion, separately, to the standard covariates. The association was no longer significant when controlling CA1 (r = −0.029, p = 0.582), CA2/3/DG (r = 0.079, p = 0.272). We next asked whether CA1 and CA2/3/DG each explained unique variance in repetition learning by entering reactivation in the other region as an additional covariate. CA1 reactivation remained a significant positive predictor after controlling for CA2/3/DG (r = 0.137, p = 0.019), whereas CA2/3/DG reactivation did not predict learning after controlling for CA1 (r = −0.003, p = 0.953). All p values are FDR-corrected. Together, these results indicate that CA1 makes a unique contribution to repetition learning beyond CA2/3/DG.

### CA1 one-step forward reactivation mediates the association between initial memory and repetition learning effects

To test whether one-step forward reactivation accounted for the association between initial memory and repetition learning, we fitted a simple mediation model for each ROI. Initial memory was the predictor (X), forward reactivation was the mediator (M), and repetition learning was the outcome (Y), with all paths adjusted for the standard covariates. Higher initial memory predicted weaker forward reactivation in all three ROIs (a path: CA1, β = −0.17, 95% CI [−0.27, −0.06]; CA2/3/DG, β = −0.14 [−0.24, −0.04]; HIPP, β = −0.14 [−0.24, −0.03]; ps ≤ .013). With initial memory controlled, reactivation predicted greater repetition learning in CA1 (b path: β = 0.07 [0.02, 0.14], p = .021) and HIPP (β = 0.06 [0.01, 0.12], p = .044), but not in CA2/3/DG (β = 0.04 [−0.01, 0.11], p = .12). Accordingly, the indirect effect was significant for CA1 (a × b = −0.012, 95% CI [−0.031, −0.002], p = .021) and HIPP (a × b = −0.009 [−0.026, −0.001], p = .036), but not for CA2/3/DG (a × b = −0.006 [−0.021, 0.001], p = .082; **Fig. 6**). All p values were FDR-corrected. Participants with lower initial memory thus showed stronger CA1 and hippocampal forward reactivation, which in turn predicted greater gains from repetition. We also repeated the analysis with short-term memory after repetition (Rep2) as the outcome. This control analysis yielded the same significant indirect effects in CA1 and HIPP (**Supplementary Results**).

**Fig. 6.**
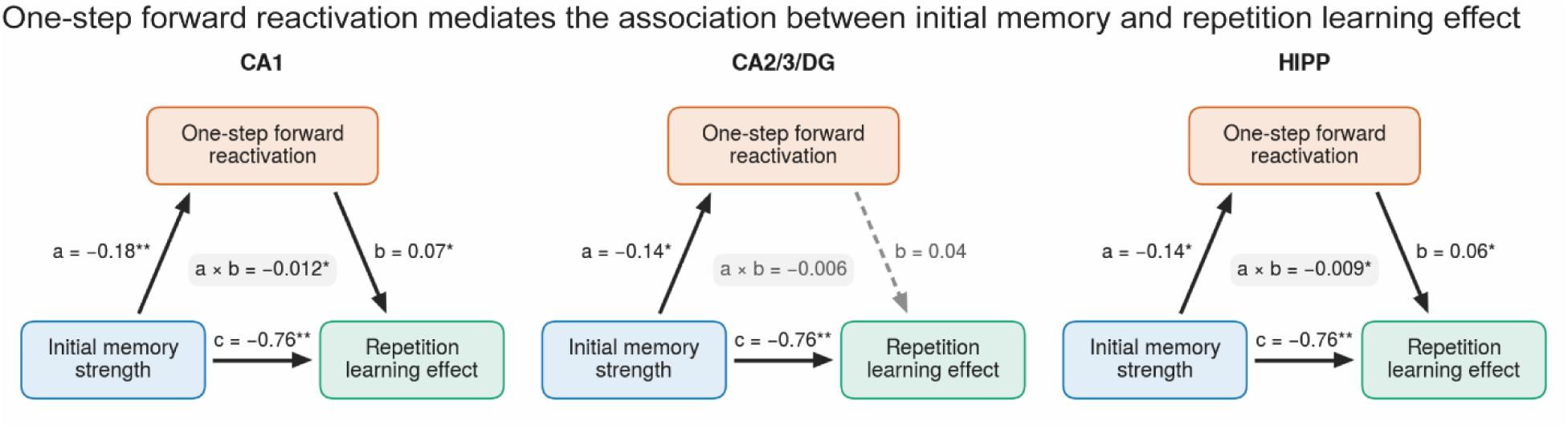
One-step forward reactivation in CA1 and HIPP mediate the association between initial memory and the repetition learning effect. In each diagram, path a is the effect of initial memory on one-step forward reactivation, path b is the effect of reactivation on the repetition learning effect after controlling for initial memory, and c is the total effect. All coefficients are standardized and adjusted for age, sex, mean head motion, education history, scanner site and normalized regional volume. The indirect effect, the product of paths a and b, is reported with its 95% confidence interval from a percentile bootstrap with 10,000 resamples, all p values were FDR-corrected. *** p < 0.001; ** p < 0.01; * p < 0.05. All p values were FDR-corrected.

Additional control analyses using overall forward reactivation as the mediator, initial memory as the predictor, and repetition learning as the outcome revealed no significant indirect effects in any ROI (ps = 0.81, FDR-corrected).

### Primacy and recency effects in backward and forward reactivation

We next tested whether our findings were driven by serial position biases, such as primacy or recency effects, by examining whether repetition learning effects were associated with increased backward reactivation of the first sequence item or forward reactivation of the final item during Rep2 encoding.

We did not find significant relationships for either forward or backward reactivation after accounting for standard covariates (all ps > 0.435, FDR-corrected).

### Forward reactivation is independent of working memory capacity

Given the sequential nature of the face-name associative task, an alternative explanation is that the robust forward reactivation observed during Rep2 encoding merely reflects active working memory (WM) maintenance rather than experience-dependent updating of episodic memory representations. To test this, we related individuals’ working memory capacity, measured by an out-of-scanner List Sorting Working Memory Test, to hippocampal forward and backward reactivation.

Individual differences in working memory capacity were not significantly associated with either overall forward reactivation or one-step forward reactivation during Rep2 encoding across any ROI after adjusting for standard covariates (all ps > 0.675) .

## Robustness with respect to hippocampal template

To demonstrate that our findings are robust with respect to the choice of hippocampal atlas, we re-segmented the MTL using the ASHS-Princeton Young Adult atlas, conducted a series of control analyses, and replicated all the main findings (see **Supplementary Results**).

## Discussion

The robust behavioral benefits of repetition learning have long been recognized as a critical mechanism for preserving episodic memory across the adult lifespan. However, the precise neural computations underpinning these gains have remained poorly understood. Our findings provide a novel computational framework by demonstrating four key principles of repetition learning: (1) experience-dependent changes in hippocampal representations from initial to repeated encoding predict behavioral memory improvements; (2) these improvements are driven by active, anticipatory forward reactivation rather than retrospective backward reactivation, even when such prediction is entirely incidental to task demands; (3) forward reactivation serves an adaptive, compensatory function that prioritizes initially weak memories; and (4) this precise, predictive mechanism underlying repetition learning in CA1 remains robust after adjustment for age, regional volume, education, sex, head motion, scanner site, backward reactivation, initial memory strength, and contribution from CA2/3/DG and other MTL subfields, and cannot be explained by temporal carryover, serial-position bias, stimulus order, or general working-memory capacity. Together, these findings characterize repetition learning as an active, anticipatory process supported by a hippocampal mechanism that is stable and resilient across aging.

### The hippocampus as a predictive engine for repetition learning

A central theoretical advance of the current study is the demonstration that the mnemonic benefits of repeated exposure are accompanied by active, anticipatory forward prediction. Traditional study-phase retrieval models posit that repetition learning relies on the online reinstatement of previously encoded traces ^11^. Our findings extend this account by showing that reactivation during repeated encoding is not confined to reinstating the past but is directed toward the immediate future. Experience-dependent representational changes in CA1, CA2/3/DG, and the whole hippocampus tracked repetition-related memory gains, and directional decomposition of these changes revealed a selective contribution of forward reactivation of upcoming sequence items rather than backward reactivation of preceding items. Forward reactivation accounted for unique variance in memory improvement beyond backward reactivation, and this relationship was temporally precise, emerging selectively for the immediately upcoming item. Repetition may therefore strengthen memory not simply by refreshing previously encoded traces, but by transforming prior experience into structured expectations about what will occur next.

Several features support this prospective interpretation. The forward-reactivation effect was sequence-specific and could not be readily attributed to serial-position biases, stimulus location/order, cross-repetition same-item similarity, or nonspecific temporal carryover. Moreover, forward prediction was not explicitly required by the task, suggesting that the hippocampus spontaneously exploited the learned transition structure during re-encoding. Forward reactivation was also unrelated to individual differences in working-memory capacity, arguing against the possibility that it merely reflected active maintenance and instead supporting its interpretation as experience-dependent updating of episodic representations. Together, these dissociations extend the traditional emphasis on the hippocampus as a primarily retrospective memory system. As sequence learning progresses, hippocampal representations may operate as a predictive map, leveraging prior encoding to anticipate the immediately upcoming state ^35,36,56^.

Our findings highlight CA1 as a candidate substrate linking temporally organized memory representations to the benefits of repeated encoding. Although one-step forward reactivation predicted repetition-related gains in both CA1 and CA2/3/DG as well as the whole hippocampus, only CA1 explained unique variance after accounting for reactivation in the other MTL regions. Its association with repetition learning also survived adjustment for initial memory strength in a separate mediation analysis. This contribution is consistent with converging evidence for CA1 involvement in temporal organization, prediction, and memory integration. Rodent studies show that CA1 time cells encode successive moments within experiences and that gradually evolving ensemble activity supports memory for event order ^57–59^. Complementary human studies show that hippocampal representations bind items to their positions within learned sequences ^60^, while greater representational similarity during encoding predicts both remembered temporal proximity ^61^ and successful temporal-order memory ^62^. Such temporal organization provides a potential basis for a familiar item to cue its learned successor during re-exposure. CA1’s convergence of inputs from CA3 and entorhinal cortex further positions it to relate retrieved information to ongoing experience, consistent with its sensitivity to violations of learned temporal expectations ^63^.

Computational models implicate the entorhinal–CA1 pathway in learning regularities across experiences ^64^, while human imaging studies associate CA1 with integration of related memories during encoding ^65,66^. Together, these findings suggest that anticipatory reinstatement may prepare an upcoming representation for renewed encoding, allowing learned temporal relationships to guide memory strengthening. Our results extend this framework by linking one-step forward reactivation during intact sequence repetition to associative memory gains, with CA1 contributing information beyond the signals measured in other MTL regions. This pattern supports a prospective role for CA1 temporal representations in repetition learning, although the underlying cellular mechanisms and interactions among subfields remain unresolved.

### Adaptive prioritization of weakly encoded memories

Crucially, we observed that this forward predictive mechanism is highly adaptive: forward prediction during the second encoding phase was negatively associated with short-term memory performance during the initial exposure across the CA1, CA2/3/DG, and the whole hippocampus. This indicates that initially weakly encoded items selectively trigger the strongest anticipatory forecasting during re-exposure, successfully rescuing vulnerable memory traces and driving immediate behavioral gains.

This adaptive allocation of neural resources during active repetition elegantly aligns with, and extends, established computational frameworks regarding memory consolidation ^67^. Prominent theoretical models (e.g., Competitive Trace Theory, Non-Monotonic Plasticity) and empirical studies of offline replay have long posited that during post-encoding rest or sleep ^68–70^, the hippocampus actively prioritizes the spontaneous reactivation of weakly encoded representations to prevent their degradation ^71,72^. Our findings reveal that this prioritization principle is not restricted to offline consolidation states. Instead, it actively governs online repetition learning. When re-exposed to a sequence, the hippocampal predictive map dynamically gauges trace vulnerability, scaling the magnitude of forward prediction inversely with initial encoding strength.

### Within-sequence forward reactivation shows no detectable age dependence

While age-related episodic-memory decline has been linked to reduced fidelity, distinctiveness, and stability of neural representations, human studies have largely focused on reinstatement of individual items or encoding patterns lacking an explicit temporal structure ^47,48,73–78^, leaving how sequential reactivation changes with age comparatively unclear. Rodent studies indicate that aging alters hippocampal sharp-wave-ripple dynamics, transient population events during which recently experienced neural sequences are commonly replayed ^79^, and reduces the temporal fidelity of sequence reactivation; nevertheless, stronger sequence replay remains associated with better spatial memory ^80–82^. Similarly, a recent human EEG study reported reduced sequential reactivation of navigational events during explicit retrieval in middle-aged adults, while stronger reactivation was associated with better spatial memory ^83^. Such findings are consistent with broader evidence that neural mechanisms supporting memory can remain related to individual differences in performance irrespective of chronological age, potentially contributing to resilience in cognitive aging ^42,84,85^. Neither forward reactivation nor the behavioral benefit of repetition varied detectably with age, and their association remained robust after adjustment for age and other demographic, structural, and imaging covariates. This pattern extends evidence that hippocampal contributions to memory encompass both age-sensitive and age-independent components ^42^. Thus, although aging may compromise the fidelity or temporal organization of mnemonic representations, the prospective use of ongoing reactivation to strengthen upcoming memories may remain relatively preserved across adulthood and represent a candidate mechanism supporting resilience in cognitive aging.

### Limitations and future directions

Several limitations of the current study warrant consideration. First, the inherent limits of the BOLD signal required the functional grouping of the CA2, CA3, and DG subfields, necessitating future high-field (e.g., 7T) imaging to disentangle these discrete microcircuits ^86^. Second, our representational similarity analysis captures macro-level mnemonic states but lacks the millisecond temporal resolution required to directly observe the rapid oscillatory dynamics that govern forward prediction in neurophysiological models. Finally, utilizing a constrained set of ten unique stimuli meant our analyses necessarily relied on inter-individual differences. Future studies should employ longer sequences with higher stimulus densities to directly track these experience-dependent representational shifts within individual subjects.

## Conclusion

In summary, our findings demand a theoretical pivot in how we conceptualize the neural computations underlying repetition learning. Rather than relying on deliberate retrospective rehearsal, sequence learning is driven by a spontaneous, temporally precise forward-predictive mechanism that adaptively prioritizes vulnerable memory traces. CA1 makes a distinctive contribution to this process, predicting memory gains beyond other hippocampus subfields. The association between forward reactivation and memory improvement remained robust after accounting for age and other covariates, while neither reactivation nor the behavioral benefit of repetition varied detectably with age, suggesting that the capacity to use learned sequence structure during re-encoding is relatively preserved across mid-to-late adulthood. Together, these findings identify forward reactivation in CA1 as a distinctive neural signature of repetition learning that appears preserved across mid-to-late adulthood.

## Methods

### Dataset

We utilized the Human Connectome Project in Aging (HCP-A) dataset ^54^, which initially included 680 participants with available neuroimaging and behavioral data for the face-name task. To ensure data quality, we excluded participants based on criteria such as excessive head motion (defined as a mean frame displacement (FD) exceeding 0.15mm or a maximum frame displacement exceeding 2mm), or very poor memory performance (mean accuracy for in scanner memory less than 0.1). This screening process resulted in the exclusion of 256 participants. Additionally, a further 6 participants were excluded from the remaining sample due to suboptimal segmentation of hippocampus subfields. The final sample comprised 418 participants (mean age = 56.13 years, SD = 13.47, range = 36–100; 236 female). Participants had, on average, 17.78 years of education (SD = 2.02). Cognitive status was assessed using the Montreal Cognitive Assessment (MoCA), with scores ranging from 20 to 30 (mean = 26.87, SD = 2.26).

For details regarding ethical procedures, informed consent, and participant recruitment, please refer to the comprehensive description provided by Bookheimer and colleagues^54^.

### Face name task

The FaceName task consists of a single 276-second run, involving encoding, distractor, and recall blocks that are repeated twice for each set of faces. During the task, participants are instructed to memorize the names associated with a series of faces during the encoding blocks and subsequently attempt to recall them silently during the recall blocks. The task commences with a countdown, followed by the first encoding block lasting 22 seconds. This block includes a 2-second cue to MEMORIZE (Encoding), followed by the presentation of 5 face/name pairs shown for 4 seconds each. The distractor block ensues, comprising a 2-second cue followed by a 20-second Go/NoGo task. Subsequently, the recall block is presented, starting with a 2-second cue to RECALL, followed by a 20-second interval where the same faces are shown with “???” (without their corresponding names) for 4 seconds each. Participants are instructed to press the left button with their index finger when they see each face/name pair appear on the screen during the encoding blocks. In the recall blocks, they are instructed to press the left button when they believe they have correctly remembered the name associated with a face. The task included two distinct sequences, each comprising a different set of five face–name pairs. The encoding and recall phases were repeated twice (Repetition 1/2, Rep1/Rep2) for each sequence, with recall phases being always followed by the encoding phase for each sequence. Participants are tested again immediately after removal from the MRI scanner at the conclusion of the scan session.

The Face-name task is always the last task performed during one neuroimaging session, therefore minimizing and standardizing the retrieval interval.

### Cognitive and clinical measures

Participants completed a comprehensive clinical and neuropsychological assessment session as part of the standard Human Connectome Project-Aging (HCP-A) protocol. Global cognitive status was evaluated using the Montreal Cognitive Assessment (MoCA), a widely validated screening instrument for detecting general cognitive impairment ^87^.

To assess domain-specific cognitive performance, we utilized two core measures from the NIH Toolbox for Assessment of Neurological and Behavioral Function ^88^. Working memory capacity was measured using the NIH Toolbox List Sorting Working Memory Test, a task that requires participants to recall and sequence visually and aurally presented stimuli (e.g., animals and food items) in size order, placing demands on both information storage and executive manipulation.

### MRI image data acquisition

HCP-A provides details of parameters for brain MRI elsewhere (Bookheimer et al., 2019; Harms et al., 2018). We used multiple brain images, covering task fMRI (Face name task), structural MRI, and high-resolution hippocampal scan. For each individual, 345 frames were acquired using multiband, gradient-echo planar imaging with the following parameters: repetition time (TR), 800 ms; echo time (TE), 37 ms; flip angle, 52°; field of view, 208×208 mm; matrix, 104×104; and voxel dimensions, 2 mm isotropic. High-resolution anatomical images of the hippocampus and surrounding cortex were acquired with a T2-weighted turbo-spin echo (TSE) anatomical sequence: 30 slices; TR: 4800 ms, TE, 106 ms; flip angle, 135°; resolution: 0.39 x 0.39 x 2mm, slices oriented perpendicular to the long axis of the hippocampus. An 8 min whole brain structural image was collected using a 3D T1-weighted MPRAGE sequence: TR, 2500 ms; TE1, 1.81 ms; TE2, 3.6 ms; TE3, 5.39 ms; TE4, 7.18 ms; flip angle, 8°; spatial resolution, 0.8 mm isotropic.

### Neuroimaging data preprocessing

Image preprocessing and statistical analysis were performed using FEAT (FMRI Expert Analysis Tool) version 6.00, part of FSL (FMRIB software library, version 5.0.9, www.fmrib.ox.ac.uk/fsl). No spatial smoothing was performed. The data were filtered in the temporal domain using a nonlinear high-pass filter with a 100-s cutoff. Following Deuker et al. ^89^ and Bellmund et al. ^90^, 6 motion parameters were used as predictors in a GLM. The residuals from this model (which could not be explained by motion) were then taken into the next analysis step. All analyses were conducted in native space.

### MTL subfield demarcation and ROIs

Medial temporal lobe (MTL) subfields were segmented using Automatic Segmentation of Hippocampal Subfields (ASHS) based on each participant’s high-resolution T2-weighted MRI scans ^91^. Segmentation was performed using the ASHS-Princeton-1.0.0-OlderAdult atlas. The MTL was parcellated into CA1, CA2/3, dentate gyrus (DG), subiculum (SUB), perirhinal cortex (PRC), entorhinal cortex (ERC), and parahippocampal cortex (PHC). Due to resolution limitations, CA2/3 and DG were combined into a single subfield (CA2/3/DG).

All subfield regions of interest (ROIs) were transformed into each participant’s native space using Advanced Normalization Tools (ANTs v2.3.5). Segmentation quality was assessed via visual inspection, and 6 participants were excluded due to suboptimal segmentation. All primary analyses were based on the older-adult atlas. To ensure that findings were not dependent on atlas selection, we repeated the segmentation using a young-adult atlas (Princeton Young Adult 3T ASHS Atlas). The MTL was similarly parcellated into CA1, CA2/3/DG, SUB, PRC, ERC, and PHC. The main results were replicated and are reported in the Supplementary Materials.

Time point–level activity maps were subsequently extracted from two ROIs (CA1 and CA2/3/DG) and the whole hippocampus (HIPP) for each participant for downstream analyses.

### Multivariate pattern similarity analysis (MPS)

#### Experience-dependent memory reinstatement in relation to enhanced memory performance in repetition learning

The 5^th^ and 6^th^ TRs (4 ∼ 5.6s) after stimuli onset corresponding to the peak of the BOLD response were averaged for each trial. MPS was applied to measure the similarity of activation patterns by calculating the correlation between trials in each hippocampus subfield. We constructed a neural similarity matrix to quantify the pattern similarity between any pair of items within the same sequence and within the repetition. We then measured changes of the averaged pattern similarity values (the upper triangle elements) between repetitions. Because the perceptual property, trial structure, and the sequence were identical between repetitions, the changes in pattern similarity could be solely attributed to learning experiences. The experience-dependent pattern similarity changes were then correlated with memory performance differences between repetitions across individuals. Cross-sequence pattern similarity was quantified separately for Rep1 and Rep2 as the similarity between stimulus representations from two different sequences. As a control analysis, we also examined whether changes in cross-sequence pattern similarity were associated with repetition-related learning effects.

#### Confirming the direction of experience-dependent pattern reinstatement

Two potential mechanisms could drive these learning-related changes in hippocampal pattern similarity: (1) forward prediction, wherein participants increasingly anticipate upcoming stimuli while encoding the current item during the second repetition (Rep2); and (2) backward reactivation, wherein participants rehearse or reactivate representations of recently encountered items. To distinguish between these possibilities, we conducted a directional pattern similarity analysis. For each stimulus presented during Rep2, if allowable, we calculated its neural pattern similarity with both succeeding and preceding stimuli from the first repetition (Rep1) to generate independent indices of forward and backward reactivation, respectively.

These directional measures were then correlated with individual differences in the behavioral repetition learning effect (i.e., memory improvement from Rep1 to Rep2). A significant correlation specific to one direction would indicate that either prediction or rehearsal primarily drives the experience-dependent representational changes.

Alternatively, if both forward and backward reactivation simultaneously contribute to these representational changes, both measures should demonstrate significant relationships with the repetition learning effect. To assess whether these effects reflect memory for sequence position rather than stimulus-specific content, we conducted additional control analyses examining the relationship between cross-sequence forward and backward reactivation and repetition-related learning effects. Cross-sequence forward reactivation was quantified as the similarity between Rep2 stimulus (k) and stimulus after the order k from the other sequence Rep1; cross-sequence backward reactivation was quantified as the similarity between Rep2 stimulus (k) and stimulus preceding the order k from the other sequence Rep1.

Furthermore, to quantify the relative contribution of forward prediction versus backward reactivation, we computed a composite directional index (the difference between the forward and backward reactivation metrics). We then correlated this index with the behavioral repetition effect. A positive correlation would suggest that forward prediction dominates the brain-behavior relationship, whereas a negative correlation would indicate that backward reactivation is the primary driver.

Because all participants experienced an identical temporal structure of sequence encoding and recall, any influence of temporal distance (or lag) on pattern similarity should be comparable across individuals and thus is unlikely to drive our findings. The same logic applies to our other analyses.

#### Linking forward prediction and backward reactivation to initial short-term memory

After establishing the presence of forward and/or backward reactivation during Rep2, we investigated the behavioral factors that modulate the strength of these neural phenomena. To determine whether initial encoding success dictated subsequent neural dynamics, we examined the relationship between short-term memory performance during Rep1 and the magnitude of forward/backward reactivation during Rep2.

#### Relating the temporal distance of memory prediction to individual differences in memory improvement

To investigate the temporal extent of forward and backward reactivation, we examined whether these processes vary based on the sequential distance between items, as not all preceding or succeeding stimuli may contribute equally. Therefore, we quantified cross-repetition neural pattern similarity as a function of distance (i.e., lag) and examined how these distance-dependent metrics related to repetition learning effect.

Specifically, for a given trial (e.g., the second stimulus) during Rep2, we computed its pattern similarity with items presented at varying distance steps during Rep1. These steps ranged from immediately adjacent items (distance 1; lag +1 or −1) extending progressively to encompass more temporally distant items within the sequence. Similarity values were averaged within each discrete distance step and subsequently correlated with individual repetition learning effects to determine how the temporal horizon of prediction or rehearsal supports memory enhancement.

### Mediation analysis

To test whether one-step forward reactivation accounted for the association between initial memory and repetition learning, we fitted a mediator model for CA1, CA2/3/DG and the whole hippocampus (HIPP) using the CANlab Mediation Toolbox ^92^ in MATLAB R2022b. Initial memory was the predictor, one-step forward reactivation in the ROI the mediator, and repetition learning the outcome. Age, sex, head motion, education, scanner site and ROI volume were included as covariates in all regressions. The model estimates paths a (initial memory to reactivation) and b (reactivation to repetition learning, controlling for initial memory), the total (c) and direct (c′) effects, and the indirect effect (a × b). Continuous variables were z-scored, so paths are standardized coefficients. Significance was assessed with 10,000 bootstrap resamples of participants, using 95% bias-corrected and accelerated (BCa) confidence intervals and two-sided BCa p values ^93^, both computed with MATLAB’s ‘bootci’. We also conducted two additional mediation analyses, one with short-term memory after repetition as the outcome and one with overall forward reactivation as the mediator.

### Control analyses for serial position effects

To ensure that the observed associations between neural reactivation and behavioral memory improvement were not confounded by general serial position effects (e.g., primacy and recency biases), we conducted a targeted control analysis. Specifically, for each item presented during Rep2, we calculated its neural pattern similarity with the first and last items of the corresponding sequence. This approach allowed us to isolate and quantify the neural prediction or rehearsal specifically directed at sequence boundaries. These boundary-specific similarity values were then averaged across all valid trial pairs and correlated with individual repetition learning effects to confirm that our primary findings were not disproportionately driven by the high saliency of these sequence positions.

## Data availability

The data used in this study are publicly available from the Human Connectome Project (Connectome Coordinating Facility) and the NIMH Data Archive (NDA; Collection ID: 2847). Access to the dataset is subject to the respective data use agreements and can be obtained through the official repositories.

## Code availability

Functional MRI data preprocessing and statistical analyses were performed on the FSL6, ANTs (v2.3.5), and Matlab 2022b. Code to analyze the data is available at OSF: https://osf.io/4m92e.

## Acknowledgements

This research was supported by Alzheimer’s Association grants AARGD-NTF-21-850781 (WC), and NIH grants AG093006 (WC), NS086085 and MH126518 (VM), the Wu Tsai Neurosciences Institute at Stanford University (VM), and Arizona Alzheimer’s Consortium & State of Arizona DHS Pilot Grant (L.Z.).

## References

1 Ebbinghaus, H. A contribution to experimental psychology. New York, NY: Teachers College, Columbia University (1913).

2 Brown, P. C., Roediger III, H. L. & McDaniel, M. A. in Make It Stick (Harvard University Press, 2015).

3 Karpicke, J. D. & Roediger III, H. L. The critical importance of retrieval for learning. science 319, 966–968 (2008).

4 Hintzman, D. L. Repetition and memory. Psychology of learning and motivation 10, 47–91 (1976).

5 Kim, K., Hsieh, L.-T., Parvizi, J. & Ranganath, C. Neural repetition suppression effects in the human hippocampus. Neurobiology of Learning and Memory 173, 107269 (2020).

6 Vilberg, K. L. & Davachi, L. Perirhinal-hippocampal connectivity during reactivation is a marker for object-based memory consolidation. Neuron 79, 1232–1242 (2013).

7 Reagh, Z. M., Murray, E. A. & Yassa, M. A. Repetition reveals ups and downs of hippocampal, thalamic, and neocortical engagement during mnemonic decisions. Hippocampus 27, 169–183 (2017).

8 Xiao, X., Li, J., Cao, D., Zhang, J. & Jiang, T. Contributions of repeated learning to memory in humans: insights from single-neuron recordings in the hippocampus and amygdala. Cerebral Cortex 34, bhae244 (2024).

9 Norman, K. A., Polyn, S. M., Detre, G. J. & Haxby, J. V. Beyond mind-reading: multi-voxel pattern analysis of fMRI data. Trends in cognitive sciences 10, 424–430 (2006).

10 Rissman, J. & Wagner, A. D. Distributed representations in memory: insights from functional brain imaging. Annual review of psychology 63, 101–128 (2012).

11 Halpern, D. J. et al. Study-phase reinstatement predicts subsequent recall. Nature neuroscience 28, 883–890 (2025).

12 Thios, S. J. & D’Agostino, P. R. Effects of repetition as a function of study-phase retrieval. Journal of Verbal Learning and Verbal Behavior 15, 529–536 (1976).

13 Hintzman, D. L., Summers, J. J. & Block, R. A. Spacing judgments as an index of study-phase retrieval. Journal of Experimental Psychology: Human Learning and Memory 1, 31 (1975).

14 Hu, X., Cheng, L. Y., Chiu, M. H. & Paller, K. A. Promoting memory consolidation during sleep: A meta-analysis of targeted memory reactivation. Psychological bulletin 146, 218 (2020).

15 Rasch, B. & Born, J. Maintaining memories by reactivation. Current opinion in neurobiology 17, 698–703 (2007).

16 Schlichting, M. L. & Preston, A. R. Memory reactivation during rest supports upcoming learning of related content. Proceedings of the National Academy of Sciences 111, 15845–15850 (2014).

17 Tambini, A. & Davachi, L. Persistence of hippocampal multivoxel patterns into postencoding rest is related to memory. Proceedings of the National Academy of Sciences 110, 19591–19596 (2013).

18 Wamsley, E. J. Offline memory consolidation during waking rest. Nature Reviews Psychology 1, 441–453 (2022).

19 Liu, Y., Nour, M. M., Schuck, N. W., Behrens, T. E. & Dolan, R. J. Decoding cognition from spontaneous neural activity. Nature Reviews Neuroscience 23, 204–214 (2022).

20 Chen, Z. et al. Interpreting human sleep activity through neural contrastive learning. Neuron (2026).

21 Silva, M., Baldassano, C. & Fuentemilla, L. Rapid memory reactivation at movie event boundaries promotes episodic encoding. Journal of Neuroscience 39, 8538–8548 (2019).

22 Chen, Z. S. & Wilson, M. A. How our understanding of memory replay evolves. Journal of Neurophysiology 129, 552–580 (2023).

23 Bubic, A., Von Cramon, D. Y. & Schubotz, R. I. Prediction, cognition and the brain. Frontiers in human neuroscience 4, 1094 (2010).

24 Bar, M. The proactive brain: memory for predictions. Philosophical Transactions of the Royal Society B: Biological Sciences 364, 1235–1243 (2009).

25 Aly, M., Barense, M. & Kok, P. Vol. 381 20250236 (The Royal Society, 2026).

26 Ólafsdóttir, H. F., Bush, D. & Barry, C. The role of hippocampal replay in memory and planning. Current Biology 28, R37–R50 (2018).

27 Diba, K. & Buzsáki, G. Forward and reverse hippocampal place-cell sequences during ripples. Nature neuroscience 10, 1241–1242 (2007).

28 Foster, D. J. Replay comes of age. Annual review of neuroscience 40, 581–602 (2017).

29 Kahana, M. J. Associative retrieval processes in free recall. Memory & cognition 24, 103–109 (1996).

30 Howard, M. W., Fotedar, M. S., Datey, A. V. & Hasselmo, M. E. The temporal context model in spatial navigation and relational learning: toward a common explanation of medial temporal lobe function across domains. Psychological review 112, 75 (2005).

31 Drosopoulos, S., Windau, E., Wagner, U. & Born, J. Sleep enforces the temporal order in memory. PLoS One 2, e376 (2007).

32 Paz, R. et al. A neural substrate in the human hippocampus for linking successive events. Proceedings of the National Academy of Sciences 107, 6046–6051 (2010).

33 Schapiro, A. C., Kustner, L. V. & Turk-Browne, N. B. Shaping of object representations in the human medial temporal lobe based on temporal regularities. Current biology 22, 1622–1627 (2012).

34 O’Reilly, R. C. & Norman, K. A. Hippocampal and neocortical contributions to memory: Advances in the complementary learning systems framework. Trends in cognitive sciences 6, 505–510 (2002).

35 Davachi, L. & DuBrow, S. How the hippocampus preserves order: the role of prediction and context. Trends in cognitive sciences 19, 92–99 (2015).

36 Stachenfeld, K. L., Botvinick, M. M. & Gershman, S. J. The hippocampus as a predictive map. Nature neuroscience 20, 1643–1653 (2017).

37 Buckner, R. L. The role of the hippocampus in prediction and imagination. Annual review of psychology 61, 27–48 (2010).

38 Barron, H. C., Auksztulewicz, R. & Friston, K. Prediction and memory: A predictive coding account. Progress in neurobiology 192, 101821 (2020).

39 Strange, B. A., Witter, M. P., Lein, E. S. & Moser, E. I. Functional organization of the hippocampal longitudinal axis. Nature reviews neuroscience 15, 655–669 (2014).

40 Poppenk, J., Evensmoen, H. R., Moscovitch, M. & Nadel, L. Long-axis specialization of the human hippocampus. Trends in cognitive sciences 17, 230–240 (2013).

41 Rolls, E. T. The mechanisms for pattern completion and pattern separation in the hippocampus. Frontiers in systems neuroscience 7, 74 (2013).

42 Zheng, L. et al. Hippocampal contributions to novel spatial learning are both age-related and age-invariant. Proceedings of the National Academy of Sciences 120, e2307884120 (2023).

43 Rönnlund, M., Nyberg, L., Bäckman, L. & Nilsson, L.-G. Stability, growth, and decline in adult life span development of declarative memory: cross-sectional and longitudinal data from a population-based study. Psychology and aging 20, 3 (2005).

44 Old, S. R. & Naveh-Benjamin, M. Differential effects of age on item and associative measures of memory: a meta-analysis. Psychology and aging 23, 104 (2008).

45 Small, S. A., Schobel, S. A., Buxton, R. B., Witter, M. P. & Barnes, C. A. A pathophysiological framework of hippocampal dysfunction in ageing and disease. Nature Reviews Neuroscience 12, 585–601 (2011).

46 Raz, N. & Rodrigue, K. M. Differential aging of the brain: patterns, cognitive correlates and modifiers. Neuroscience & Biobehavioral Reviews 30, 730–748 (2006).

47 Koen, J. D. & Rugg, M. D. Neural dedifferentiation in the aging brain. Trends in cognitive sciences 23, 547–559 (2019).

48 Yassa, M. A., Mattfeld, A. T., Stark, S. M. & Stark, C. E. Age-related memory deficits linked to circuit-specific disruptions in the hippocampus. Proceedings of the National Academy of Sciences 108, 8873–8878 (2011).

49 Buchler, N. G., Faunce, P., Light, L. L., Gottfredson, N. & Reder, L. M. Effects of repetition on associative recognition in young and older adults: item and associative strengthening. Psychology and aging 26, 111 (2011).

50 Mitchell, D. B. How many memory systems? Evidence from aging. *Journal of Experimental Psychology: Learning*, Memory, and Cognition 15, 31 (1989).

51 Cohen, R. L., Sandler, S. P. & Schroeder, K. Aging and memory for words and action events: effects of item repetition and list length. Psychology and aging 2, 280 (1987).

52 Dutli, J., Oberauer, K. & Bartsch, L. M. Is repetition learning preserved in old age? Insights from the Hebb paradigm. Psychology and Aging 41, 319 (2026).

53 Kriegeskorte, N., Mur, M. & Bandettini, P. A. Representational similarity analysis-connecting the branches of systems neuroscience. Frontiers in systems neuroscience 2, 249 (2008).

54 Bookheimer, S. Y. et al. The lifespan human connectome project in aging: an overview. Neuroimage 185, 335–348 (2019).

55 Zheng, L., Gao, Z., McAvan, A. S., Isham, E. A. & Ekstrom, A. D. Partially overlapping spatial environments trigger reinstatement in hippocampus and schema representations in prefrontal cortex. Nature Communications 12, 6231 (2021).

56 Brunec, I. K. & Momennejad, I. Predictive representations in hippocampal and prefrontal hierarchies. Journal of Neuroscience 42, 299–312 (2022).

57 Eichenbaum, H. Time cells in the hippocampus: a new dimension for mapping memories. Nature Reviews Neuroscience 15, 732–744 (2014).

58 MacDonald, C. J., Lepage, K. Q., Eden, U. T. & Eichenbaum, H. Hippocampal “time cells” bridge the gap in memory for discontiguous events. Neuron 71, 737–749 (2011).

59 Manns, J. R., Howard, M. W. & Eichenbaum, H. Gradual changes in hippocampal activity support remembering the order of events. Neuron 56, 530–540 (2007).

60 Hsieh, L.-T., Gruber, M. J., Jenkins, L. J. & Ranganath, C. Hippocampal activity patterns carry information about objects in temporal context. Neuron 81, 1165–1178 (2014).

61 Ezzyat, Y. & Davachi, L. Similarity breeds proximity: pattern similarity within and across contexts is related to later mnemonic judgments of temporal proximity. Neuron 81, 1179–1189 (2014).

62 DuBrow, S. & Davachi, L. Temporal memory is shaped by encoding stability and intervening item reactivation. The Journal of Neuroscience 34, 13998–14005 (2014).

63 Chen, J., Cook, P. A. & Wagner, A. D. Prediction strength modulates responses in human area CA1 to sequence violations. Journal of Neurophysiology 114, 1227–1238 (2015).

64 Schapiro, A. C., Turk-Browne, N. B., Botvinick, M. M. & Norman, K. A. Complementary learning systems within the hippocampus: a neural network modelling approach to reconciling episodic memory with statistical learning. Philosophical Transactions of the Royal Society B: Biological Sciences 372, 20160049 (2017).

65 Molitor, R. J., Sherrill, K. R., Morton, N. W., Miller, A. A. & Preston, A. R. Memory reactivation during learning simultaneously promotes dentate gyrus/CA2, 3 pattern differentiation and CA1 memory integration. The Journal of Neuroscience 41, 726–738 (2021).

66 Schlichting, M. L., Zeithamova, D. & Preston, A. R. CA1 subfield contributions to memory integration and inference. Hippocampus 24, 1248–1260 (2014).

67 McClelland, J. L., McNaughton, B. L. & O’Reilly, R. C. Why there are complementary learning systems in the hippocampus and neocortex: insights from the successes and failures of connectionist models of learning and memory. Psychological review 102, 419 (1995).

68 Tambini, A. & Davachi, L. Awake reactivation of prior experiences consolidates memories and biases cognition. Trends in cognitive sciences 23, 876–890 (2019).

69 Bendor, D. & Wilson, M. A. Biasing the content of hippocampal replay during sleep. Nature neuroscience 15, 1439–1444 (2012).

70 Brodt, S., Inostroza, M., Niethard, N. & Born, J. Sleep—A brain-state serving systems memory consolidation. Neuron 111, 1050–1075 (2023).

71 Schapiro, A. C., McDevitt, E. A., Rogers, T. T., Mednick, S. C. & Norman, K. A. Human hippocampal replay during rest prioritizes weakly learned information and predicts memory performance. Nature communications 9, 3920 (2018).

72 Cowan, E. T., Schapiro, A. C., Dunsmoor, J. E. & Murty, V. P. Memory consolidation as an adaptive process. Psychonomic Bulletin & Review 28, 1796–1810 (2021).

73 Koen, J. D., Srokova, S. & Rugg, M. D. Age-related neural dedifferentiation and cognition. Current opinion in behavioral sciences 32, 7–14 (2020).

74 Leal, S. L. & Yassa, M. A. Integrating new findings and examining clinical applications of pattern separation. Nature neuroscience 21, 163–173 (2018).

75 Zheng, L. et al. Reduced fidelity of neural representation underlies episodic memory decline in normal aging. Cerebral Cortex 28, 2283–2296 (2018).

76 Srokova, S., Aktas, A. N., Koen, J. D. & Rugg, M. D. Dissociative effects of age on neural differentiation at the category and item levels. The Journal of Neuroscience 44, e0959232023 (2024).

77 Hill, P. F., King, D. R. & Rugg, M. D. Age differences in retrieval-related reinstatement reflect age-related dedifferentiation at encoding. Cerebral Cortex 31, 106–122 (2021).

78 Pauley, C. & Sander, M. C. The role of age differences in neural representations for memory performance. Neuroscience & Biobehavioral Reviews 176, 106262 (2025).

79 Roumis, D. K. & Frank, L. M. Hippocampal sharp-wave ripples in waking and sleeping states. Current opinion in neurobiology 35, 6–12 (2015).

80 Gerrard, J. L., Burke, S. N., McNaughton, B. L. & Barnes, C. A. Sequence reactivation in the hippocampus is impaired in aged rats. The Journal of Neuroscience 28, 7883–7890 (2008).

81 Wiegand, J.-P. L. et al. Age is associated with reduced sharp-wave ripple frequency and altered patterns of neuronal variability. The Journal of Neuroscience 36, 5650–5660 (2016).

82 Cowen, S. L., Gray, D. T., Wiegand, J. P. L., Schimanski, L. A. & Barnes, C. A. Age-associated changes in waking hippocampal sharp-wave ripples. Hippocampus 30, 28–38 (2020).

83 Lee, J., Park, S.-E. & Lee, S. A. Neural reinstatement and sequential reactivation of navigational episodic memory and its age-related decline. Neuropsychologia, 109319 (2025).

84 Cabeza, R. et al. Maintenance, reserve and compensation: the cognitive neuroscience of healthy ageing. Nature reviews neuroscience 19, 701–710 (2018).

85 Stern, Y. et al. Mechanisms underlying resilience in ageing. Nature Reviews Neuroscience 20, 246–246 (2019).

86 Berron, D. et al. Strong evidence for pattern separation in human dentate gyrus. Journal of Neuroscience 36, 7569–7579 (2016).

87 Nasreddine, Z. S. et al. The Montreal Cognitive Assessment, MoCA: a brief screening tool for mild cognitive impairment. Journal of the American Geriatrics Society 53, 695–699 (2005).

88 Gershon, R. C. et al. NIH toolbox for assessment of neurological and behavioral function. Neurology 80, S2–S6 (2013).

89 Deuker, L., Bellmund, J. L., Navarro Schröder, T. & Doeller, C. F. An event map of memory space in the hippocampus. elife 5, e16534 (2016).

90 Bellmund, J. L., Deuker, L. & Doeller, C. F. Mapping sequence structure in the human lateral entorhinal cortex. elife 8, e45333 (2019).

91 Yushkevich, P. A. et al. Automated volumetry and regional thickness analysis of hippocampal subfields and medial temporal cortical structures in mild cognitive impairment. Human brain mapping 36, 258–287 (2015).

92 Wager, T. D., Davidson, M. L., Hughes, B. L., Lindquist, M. A. & Ochsner, K. N. Prefrontal-subcortical pathways mediating successful emotion regulation. Neuron 59, 1037–1050 (2008).

93 Efron, B. Better bootstrap confidence intervals. Journal of the American statistical Association 82, 171–185 (1987).

