## Supplementary material for "Forward reactivation in human hippocampal CA1 supports repetition learning and is preserved in the aging brain": SI

**Supplementary Materials**

**Supplementary Methods**

**Mediation analysis.**

We refitted the mediation model using short-term memory after Rep2 as the outcome. The predictor (initial memory), mediator (one-step forward reactivation in CA1, CA2/3/DG, or the whole hippocampus), and covariates (age, sex, head motion, education, scanner site, and ROI volume) were the same as those used in the mediation analysis reported in the main text. All continuous variables were z-scored, so all paths are standardized coefficients. Path coefficients were estimated with the CANlab Mediation Toolbox ^1^. The 95% confidence intervals and two-sided p values were obtained from 10,000 bias-corrected and accelerated (BCa) bootstrap resamples of participants ^2^. P values were FDR-corrected across the three ROIs.

**Supplementary Results**

**MTL subfield volume in relation to memory**

We assessed the relationship between normalized volume and memory performance. Significant positive association was found between in-scanner short-term memory after the second repetition (Rep2) and normalized regional volume in CA1 (r = 0.124, p = 0.037), as well as between long-term memory measured outside the scanner and normalized regional volume across all examined ROIs (r = 0.112~0.196, ps < 0.023, FDR-corrected). However, these associations did not remain significant after adjusting for age, sex, and education (all ps > 0.725, FDR-corrected). Normalized volume showed trend-level associations with the repetition learning effect in CA1, CA2/3/DG, and the whole hippocampus (HIPP) (r = 0.109~0.123, ps = 0.063, FDR-corrected) after controlling for age, sex, and education.

**Hippocampal representation change predicts learning effects**

We examined whether within-sequence neural pattern similarity was greater during Rep2 than Rep1, and whether such repetition-related changes extended beyond the Rep2-versus-Rep1 encoding comparison reported in the main text to other task phases. We further tested whether these changes were associated with repetition-related memory gains. We found similar positive correlations across other learning phases. After adjusting for confounding factors, comparing Rep2 encoding with Rep1 recall yielded significant positive effects in CA1 (r = 0.158, p = 0.008) and the whole HIPP (r = 0.117, p = 0.011), alongside a non-significant effect in CA2/3/DG (r = 0.047, p = 0.371). Similarly, comparing Rep2 recall with Rep1 encoding revealed significant positive correlations across all examined regions: CA1 (r = 0.114, p = 0.038), CA2/3/DG (r = 0.109, p = 0.038), and the whole HIPP (r = 0.121, p = 0.038; all p-values FDR-corrected). In contrast, no significant associations were detected for comparisons between Rep2 recall and Rep1 recall, or between Rep2 recall and Rep2 encoding (all ps > 0.283, FDR-corrected).

We conducted additional control analyses to determine whether representational changes related to repetition learning generalized across sequences or were sequence specific. For each repetition, we calculated pattern similarity between different sequences and averaged the resulting cross-sequence similarity estimates. We then computed the change in cross-sequence similarity from the first to the second repetition and tested its association with the repetition learning effect, controlling for age, sex, education, normalized regional volume, head motion, and scanner site. No significant associations were observed in any ROI after FDR correction (all ps ≥ .732). These null cross-sequence associations, together with the primary within-sequence findings, suggest that representational changes were specific to the repeated sequence rather than reflecting a general change shared across sequences.

**Forward reactivation in the hippocampus predicts repeated learning effects**

We reported forward reactivation during Rep2 encoding defined by pattern similarity to Rep1 encoding predicted repetition learning effects in the main text. Similar patterns were observed when comparing neural activation pattern during Rep2 encoding with both Rep1 encoding and Rep1 recall, as well as when comparing Rep2 encoding with preceding or following stimuli in the same sequence across all other phases (Rep1 encoding, Rep1 recall, and Rep2 recall). After controlling for potential confounds, we found positive associations between forward reactivation and repetition effects in CA1, CA2/3/DG, and the whole HIPP (Rep2 encoding vs. Rep1 encoding + recall: r = 0.204/0.096/0.138, p = 0.0003/0.070/0.013; Rep2 encoding vs. all other phases: r = 0.218/0.148/0.153, p = 0.0001/0.005/0.005; FDR-corrected). In contrast, after adjusting for confounding factors, no significant relationships were observed between backward reactivation and repetition effects in these task phases (all ps > 0.206, FDR-corrected).

To rule out the possibility that participants were merely memorizing the location or position within a sequence rather than the specific content of the face-name pairs, we conducted a control analysis. Specifically, we computed the forward and backward reactivation using pattern similarity pairs from different sequences. For example, for P3 in Rep2 from sequence 1 (Seq1), we calculated its pattern similarity with P4/P5 in Rep1 for forward reactivation and with P1/P2 in Rep1 for backward reactivation from sequence 2 (Seq2). If memory reactivation is content-specific, we would not expect significant brain-behavior relationships using these cross-sequence comparisons. Consistent with our hypothesis, no significant correlations emerged when computing memory reactivation using pairs from different sequences, either before or after controlling for potential confounds, in both forward and backward directions (all ps > 0.304, FDR-corrected).

We also computed a directionality index to quantify the relative strength of forward versus backward reactivation by taking their difference while controlling trial lags. For example, for P3 in Rep2, we calculated the difference between its forward prediction and backward reactivation with corresponding pairs matched on lag (i.e., P4 vs. P2 and P5 vs. P1). We then examined its relationship with the repetition learning effect across individuals. After accounting for potential confounds, a positive correlation was observed in CA1 when comparing Rep2 encoding with all other phases (r = 0.132, p = 0.036, FDR-corrected), indicating that a greater bias toward forward (vs. backward) reactivation was associated with greater memory improvement across repetitions.

**Mediation analysis with short-term memory after repetition as the outcome**

With short-term memory after Rep2 encoding as the outcome, higher initial memory again predicted weaker one-step forward reactivation in all three ROIs (the a path is identical to the main analysis). Controlling for initial memory, stronger reactivation predicted better short-term memory after Rep2 in CA1 (β = 0.09, 95% CI [0.02, 0.17], p = .021) and HIPP (β = 0.08 [0.01, 0.16], p = .044), but not in CA2/3/DG (β = 0.05 [−0.01, 0.13], p = .12). The indirect effect was significant for CA1 (a × b = −0.015, 95% BCa CI [−0.039, −0.003], p = .021) and HIPP (a × b = −0.011 [−0.032, −0.001], p = .036), but not for CA2/3/DG (a × b = −0.007 [−0.026, 0.001], p = .082). Initial memory was strongly and positively related to short-term memory after repetition (total effect c = 0.60 [0.46, 0.81] for all three ROIs; direct effect c′ = 0.60 to 0.61; all p < .001). All p values were FDR-corrected.

**Replicating the main findings using a young-adult atlas**

**Experience-dependent hippocampal representation changes predict repetition learning effects (Young-adult atlas)**

After adjusting for age, sex, years of education, regional volume, head motion, and scanner sites, we observed significant positive correlations between repetition-related changes (Rep2 encoding – Rep1 encoding) in pattern similarity and the repetition learning effect in CA1, CA2/3/DG, and HIPP (r = 0.116/0.124/0.135, ps = 0.027; FDR-corrected), indicating that experience-dependent changes in neural representations robustly predict memory improvement. We also found similar positive correlations across other learning phases. After adjusting for confounding factors, comparing Rep2 encoding with Rep1 recall yielded significant positive effects in CA1 (r = 0.154, p = 0.0097), alongside a marginal effect in HIPP (r = 0.102, p = 0.079); all p values were FDR corrected.

These findings confirmed that the relationship between hippocampal representation changes and repetition learning effects was robust and not dependent on the segmentation atlas we were using.

**Directional analysis of memory reactivation: confirming forward prediction in the hippocampus (Young-adult atlas)**

There was a significant positive relationship between forward reactivation during Rep2 encoding relative to Rep1 encoding and repetition effect in the CA2/3/DG, and the whole HIPP (CA2/3/DG: r = 0.14, p = 0.015; HIPP, r = 0.111, p = 0.049; FDR-corrected). The effects remained significant after adjusting for confounding factors such as age, sex, education, regional volume, head motion, and scanner sites in CA2/3/DG (r = 0.163, p = 0.018), with a trend level effect in HIPP (r = 0.103, p = 0.081); all p values were FDR-corrected. In contrast, no significant relationships were observed for backward reactivation/rehearsal (ps > 0.957, FDR-corrected). We also observed similar effects when comparing neural activation pattern during Rep2 encoding with both Rep1 encoding and Rep1 recall, as well as when comparing Rep2 encoding with preceding or following stimuli in the same sequence across all other phases (Rep1 encoding, Rep1 recall, and Rep2 recall). After controlling potential confounds, we found positive associations between forward reactivation and repetition effects in CA1, CA2/3/DG, and the whole HIPP (Rep2 encoding vs. Rep1 encoding + recall: r = 0.088/0.15/0.122, p = 0.096/0.013/0.031; Rep2 encoding vs. all other phases: r = 0.124/0.151/0.123, p = 0.02/0.013/0.02; FDR-corrected). In contrast, after adjusting confounding factors, no significant relationships were observed between backward reactivation and repetition effects in these task phases (all ps > 0.334, FDR-corrected).

Together, these findings provide consistent evidence that forward, but not backward, hippocampal reactivation predicts memory improvement during repeated learning, and this effect was atlas-invariant.

**Hippocampal forward reactivation in relation to short-term memory (Young-adult atlas)**

Short-term memory performance during Rep1 was negatively associated with forward reactivation during Rep2 encoding defined by the similarity to neural activation patterns during Rep1 encoding in CA23/DG (r = −0.151, p = 0.018; FDR-corrected) and showed a marginal effect in HIPP (r = -0.107, p = 0.066; FDR corrected), after controlling for standard covariates including age, sex, education, normalized volume, head motion, and scanner site. Similar effects were observed in CA2/3/DG and HIPP when forward reactivation during Rep2 encoding **was** defined by its similarity to neural activation patterns during Rep1 encoding and recall (r = −0.126/−0.115, p = 0.044/0.044; FDR-corrected) and when patterns from Rep2 encoding were compared with neural activation patterns during all other phases, after accounting for standard confounding factors (r = −0.194/−0.145, p = 0.0007/0.009; FDR-corrected).

**Working memory capacity dissociates from experience-dependent forward prediction (Young-adult atlas)**

Working memory capacity significantly modulated backward reactivation. We observed a robust inverse relationship between working memory performance and backward reactivation during Rep2 encoding. After adjusting for standard covariates, this negative relationship was significant when pattern similarity was assessed relative to Rep1 encoding (CA1 and HIPP: r = -0.144/-0.173, p = 0.014/0.005; FDR-corrected), both Rep1 encoding and recall (CA1 and HIPP: r = -0.162/-0.172, ps = 0.005; FDR-corrected).

In contrast, individual differences in working memory capacity could not explain the protective repetition learning effects driven by forward reactivation. In fact, after accounting for standard covariates, we found almost no significant associations between hippocampal representations—including either within- or cross-sequence forward reactivation—and WM capacity across any ROIs (ps > 0.55, FDR-corrected). This dissociation demonstrates that the predictive hippocampal mechanisms supporting sequence relearning operate independently of general working memory resources.

**Extent of predictive reactivation: evaluating short- to long-distance forecasts (Young-adult atlas)**

We observed significant positive associations between one-step forward reactivation (i.e., prediction of the immediately upcoming item) during Rep2 encoding and repetition-related learning effects in CA1, CA2/3/DG, and the whole HIPP (r = 0.135/0.192/0.177; p = 0.01/0.0008/0.001; FDR-corrected), after controlling for standard covariates. In contrast, repetition-related learning effects were not significantly associated with one-step backward reactivation or with reactivation at any other forward or backward distance (ps > 0.243, FDR corrected).

We also tested whether our findings were driven by serial position biases, such as primacy or recency effects, by examining whether repetition learning effects were associated with increased backward reactivation of the first sequence item or forward reactivation of the final item during Rep2 encoding. We did not find significant relationships for either forward or backward reactivation after accounting for standard covariates (ps > 0.153, FDR-corrected).

**References**

1	Wager, T. D., Davidson, M. L., Hughes, B. L., Lindquist, M. A. & Ochsner, K. N. Prefrontal-subcortical pathways mediating successful emotion regulation. *Neuron* **59**, 1037–1050 (2008).

2	Tibshirani, R. J. & Efron, B. An introduction to the bootstrap. *Monographs on statistics and applied probability* **57**, 1–436 (1993).
